# Uncovering the underlying mechanisms and whole-brain dynamics of therapeutic deep brain stimulation for Parkinson’s disease

**DOI:** 10.1101/083162

**Authors:** Victor M. Saenger, Joshua Kahan, Tom Foltynie, Karl Friston, Tipu Z. Aziz, Alexander L. Green, Tim J. van Hartevelt, Angus B. A. Stevner, Henrique M. Fernandes, Laura Mancini, John Thornton, Tarek Yousry, Patricia Limousin, Ludvic Zrinzo, Marwan Hariz, Morten L. Kringelbach, Gustavo Deco

## Abstract

Deep brain stimulation (DBS) for Parkinson’s disease is a highly effective treatment in controlling otherwise debilitating symptoms yet the underlying brain mechanisms are currently not well understood. We used whole-brain computational modeling to disclose the effects of DBS ON and OFF during collection of resting state fMRI in ten Parkinson’s Disease patients. Specifically, we explored the local and global impact of DBS in creating asynchronous, stable or critical oscillatory conditions using a supercritical bifurcation model. We found that DBS shifts the global brain dynamics of patients nearer to that of healthy people by significantly changing the bifurcation parameters in brain regions implicated in Parkinson’s Disease. We also found higher communicability and coherence brain measures during DBS ON compared to DBS OFF. Finally, by modeling stimulation we identified possible novel DBS targets. These results offer important insights into the underlying effects of DBS, which may in time offer a route to more efficacious treatments.

## Introduction

Deep Brain Stimulation (DBS) is a remarkably effective treatment for a number of otherwise treatment-resistant disorders including tremor, dystonia, and Parkinson’s disease (Bronstein et al. 2011; Kringelbach and Aziz 2011; Little et al. 2013; Lozano and Lipsman 2013; Miocinovic et al. 2013). Initially, targets for Parkinson’s Disease were discovered using the highly successful 1-methyl-4-phenyl-1,2,3,6-tetrahydropyridine (MPTP) model in higher primates (Langston et al. 1983), which helped identify a number of efficacious DBS targets and most importantly the subthalamic nucleus STN (Bergman et al. 1990; Aziz et al. 1991). Perhaps surprisingly, though, the underlying mechanisms of DBS are not yet resolved despite the fact that DBS in the STN has now helped over 150,000 patients. Initially it was thought that, similar to surgical lesions, DBS acted on local circuitry but careful analysis of the biophysical properties of the brain (Kringelbach, Jenkinson, Owen, et al. 2007) has shown the most likely mechanism of DBS is through stimulation-induced modulation of the activity of macroscopic brain networks (Vitek 2002; Kringelbach et al. 2011). Corroborating evidence has come from rodent optogenetic experiments which have shown that the therapeutic effects within the STN can be accounted for by direct selective stimulation of afferent axons projecting to this region (Gradinaru et al. 2009). Still, these studies have not resolved the nature of the whole-brain dynamics arising from DBS.

A principled approach to understanding DBS mechanisms will need to take into account the structural and functional connectivity of a given DBS target within the diseased brain and to map the ensuing changes caused by this continuous perturbation. Recent advances in computational connectomics have now produced the necessary tools to allow for careful, causal exploration of whole-brain dynamics within the underlying structural connectivity (Honey et al. 2007; Deco and Kringelbach 2014; Sporns 2014; Deco et al. 2015). Using these tools, research has demonstrated that there are specific structural “fingerprints” of structural connectivity associated with successful versus unsuccessful outcomes of DBS (Fernandes et al. 2015). In addition, the connectomic analysis of a unique dataset of pre and post-DBS diffusion tensor imaging (DTI) for Parkinson’s Disease found significant localized structural changes as a result of long-term deep brain stimulation (Van Hartevelt et al. 2014). Further, using whole-brain computational modeling on the dataset to track the ensuing changes in functional connectivity of STN DBS found Hebbian-like learning in specific STN projections (van Hartevelt et al. 2015).

Functional connectivity changes following DBS across the whole human brain were first explored using MEG in patients with DBS for chronic pain which found specific functional changes in brain activity associated with pain relief (Kringelbach, Jenkinson, Green, et al. 2007) as well as the ensuing long term changes in functional connectivity after 12 months (Mohseni et al. 2012). Subsequent studies have started to use functional MRI, having significantly reduced the risks to the patient (Boertien et al. 2011) using established safe imaging conditions (Carmichael et al. 2007; Kahan et al. 2015). A first study demonstrated a reversal in cortico-thalamic coupling during voluntary movements in Parkinson’s Disease patients with STN DBS (Kahan et al. 2012), while a follow-up study used dynamic causal modeling (DCM) of the STN network to further characterize the effective connectivity of resting state motor networks (Kahan et al. 2014). In another study, fMRI and EEG were used to track the changes following DBS of the nucleus accumbens (NAc) in patients with obsessive-compulsive disorder which was found to reduce excessive connectivity between the NAc and prefrontal cortex, with decreased frontal low-frequency oscillations during symptom provocation (Figee et al. 2013).

Taken together these studies lend strong support to the idea that therapeutic DBS works by re-balancing the brain activity of the functional and structural networks in the diseased brain (Kringelbach *et al.* 2011). Still, we are missing a mechanistic understanding of how these whole-brain networks change with DBS. In traditional thermodynamical theory, *criticality* refers to a state in which two phases are indistinguishable from one another (Mora and Bialek 2011). From this viewpoint, it has been shown that the resting brain optimally operates in a similar critical manner, at the edge of a bifurcation that represents a transition between states (Deco and Jirsa 2012). Here, we used the tools from computational connectomics to investigate the fMRI responses in ten Parkinson’s Disease patients with DBS ON and OFF compared to 16 healthy participants. This allowed us to explore the local and global impact that DBS has on resting state brain dynamics (Kringelbach et al. 2015). The advantage of using this model is that it estimates a bifurcation parameter, which locally (region-by-region) and globally describes whether a system presents asynchronous, critical or synchronous oscillations. Further, to address if turning the stimulation on immediately improves and restores global coherence and diffusion of information and helps restoring global dynamics back to a healthy state, we used several metrics that allowed the identification of global enhancements in communicability and synchrony of the network as well producing artificial local oscillatory conditions as a proxy for stimulation. In the light of earlier findings addressing large-scale changes caused by Parkinson’s Disease (van Eimeren et al. 2009; Delaveau et al. 2010; Van Hartevelt *et al.* 2014) and previous research on DBS mechanisms (Kringelbach et al. 2010), we predicted that therapeutic DBS for Parkinson’s Disease would create both global and local changes in the whole-brain dynamics.

## Materials and Methods

### Data acquisition summary

A detailed data acquisition and preprocessing description can be found in the Supplementary Material. Here we present a short description only. We studied a total of 10 patients suffering from Parkinson’s Disease (Supplementary Table 1), which received bilateral DBS in the STN for about 6 months. All patients were scanned both during active therapeutic (ON condition) and inactivated stimulation (OFF condition) to extract and compute their corresponding resting state time series. These resting state fluctuations were used to further compute their corresponding functional connectivity (FC) matrices parcellated into 90 nodes in total using the AAL template (Tzourio-Mazoyer et al. 2002) from which only the right hemisphere (Table 1) was used for the analysis to avoid artifacts arising from the connection between the electrode lead and the extension cable in the left hemisphere of the patients. Additionally, a set 16 healthy participants were recruited and scanned during rest and served as a baseline control.

**Tablel.** AAL regions and abbreviations.

| **AAL Region** | **Abbreviation** |
| --- | --- |
| Inferior temporal gyrus | InfT |
| Temporal Pole: middle temporal gyrus | TPmidT |
| Middle temporal gyrus | midT |
| Temporal Pole: superior temporal gyrus | TPSupT |
| Superior temporal gyrus | SupT |
| Heschl's gyrus | Heschl |
| Thalamus | Thal |
| Globus pallidus | Pall |
| Putamen | Put |
| Caudate nucleus | Caud |
| Paracentral lobule | Parac |
| Precuneus | Precu |
| Angular gyrus | Ang |
| Supramarginal gyrus | SupraM |
| Inferior parietal gyrus | InfP |
| Superior parietal gyrus | SupP |
| Postcentral gyrus | Postc |
| Fusiform gyrus | Fus |
| Inferior occipital gyrus | InfO |
| Middle occipital gyrus | MidO |
| Superior occipital gyrus | SupO |
| Lingual gyrus | Ling |
| Cuneus | Cun |
| Calcarine Fissure | Calc |
| Amygdala | Amyg |
| Parahippocampal gyrus | ParaHG |
| Hippocampus | Hipp |
| Posterior cingulate gyrus | PostCG |
| Middle cingulate gyrus | MidCG |
| Anterior cingulate gyrus | AntCG |
| Insula | Ins |
| Gyrus rectus | GyrR |
| Superior frontal gyrus, medial orbital | MOSupF |
| Superior frontal gyrus, medial | MSupF |
| Olfactory cortex | Olf |
| Supplementary motor area | SupplM |
| Rolandic operculum | Rolan |
| Inferior frontal gyrus, orbital | OrInfF |
| Inferior frontal gyrus, triangular | TrInfF |
| Inferior frontal gyrus, opercular | OpInfF |
| Middle frontal gyrus, orbital | OrMidF |
| Middle frontal gyrus | MidF |
| Superior frontal gyrus, orbital | OrSupF |
| Superior frontal gyrus, dorsolateral | DISupF |
| Precentral gyrus | Precen |

### Phase consistency

To assess fluctuations of functional connectivity, we evaluated the phase dispersion *φ*_*k*_(t) in the following manner. By filtering the time series with a band-pass of 0.04-0.07Hz and further using a Hilbert transform, we evaluated the instantaneous phase of every node at every time point. This allowed us to construct a matrix of phase differences, using a three-step sliding window over time. The ensuing array of phase differences was then summarized in terms of its mean and standard deviation over time. The latter characterizes fluctuations in phase synchronization and furnishes a measure of metastability (Tognoli and Kelso 2014). We repeated this process in each of the 10 participants for both the ON and OFF condition as well as in each of the 16 Healthy participants. Metrics between the ON and OFF condition were compared with a two-tailed paired t- test, while between the ON and Healthy with a two-tailed unpaired *t*-test.

### Integration and semi-random walker algorithm

The integrative features of the network were measured using two different metrics. The first one is a simple but rather powerful way to address how integrated or coupled a network is. First, a connectivity threshold *c* (goes from 0 to 1 in steps of 0.01) is gradually applied for any given FC until the matrix is fully disconnected. Then, the size of the largest component *L* in which all pair of nodes are connected is calculated for every thresholded matrix. Finally, we defined integration *I* as:

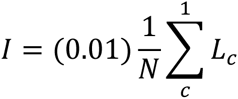

where *N* is the number of nodes in the network and *L*_*c*_ is the thresholded largest component. This was applied for all participants in both the ON and OFF conditions and compared the values with a paired *t*-test. We further calculated this same metric in the healthy set as a way to quantify a baseline control integration value and compared it to the ON condition with an unpaired *t*-test.

As a second way to measure integration under a diffusion of information flow perspective, we created a semi-random walker algorithm that measures the ability of a random walker to diffuse or explore the network. Inspired by the work done by Rosvall and Bergstrom (2007), we created a semi-random walker algorithm and measure its ability to diffuse through any FC. First, a walker is set at a random initial node in the network choosing its next node based on a probabilistic approach such that links with higher weights or correlation values are chosen more frequently. To do this, the original functional weights of each of the nodes in a given network matrix M need to be transformed in such a way that weights corresponds to out-probabilities of the random walker and the sum of all probabilities on every node is always 1. For a given weight in a node *i*, this probability is simply given by:

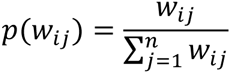

where *w_ij_* is the weight *j* of node *i* and *n* is the total number of nodes in the network. The probability matrix *p*(M) is then constructed:

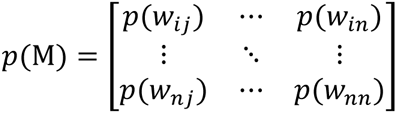

We then let the random walker explore or diffuse over *p*(M) without imposing restrictions in directionality until it has visited all nodes in the network at least one time. Then, we calculate the amount of steps *S* the walker required to diffuse over 90% of the network. This simulation is repeated 10,000 times and the mean *S* is then computed. To avoid that the random walker gets trapped too long inside a highly connected cluster, we added a fixed random teleportation probability (Brin and Page 1998) of 0.1. To check if diffusion or flow of information is affected by turning the stimulation on, this process was applied for each participant in both conditions. Diffusion of information was also calculated for the Healthy control group.

### Whole-brain Modeling

The dynamic whole-brain model uses a DTI-based structural backbone network (Van Hartevelt *et al.* 2014) of 45 brain regions (Table 1). A detailed methodological description can be found in a recently published study (Deco et al. 2016) and in the Supplementary Material. In short, this model has two sorts of parameters: a bifurcation parameter that is local to each node and a free parameter, which scales the global connectivity (coupling). Heuristically, we can consider the bifurcation parameters *a* as mediating intrinsic (or within node) dynamics, while the extrinsic (between-node) connectivity is parameterized by the global coupling *G*. In what follows, we optimized the local (bifurcation or intrinsic) parameters to ensure the relative power around each intrinsic frequency band matched the relative power observed in empirical data. This was repeated for several levels (60) of the global (extrinsic) coupling G. Having fit the parameters to empirical data, we then inferred the most likely global coupling by seeing how well it predicted a variety of functional integration measures (see below) based upon the empirical data. We then examined the intrinsic bifurcation parameters and tested for differences in their distribution over nodes (and global coupling) between the three conditions (ON, OFF and healthy). We will first describe the three metrics used to find the global coupling that best explained empirical dynamics. We then describe how their distributions were compared over conditions.

### Agreement between empirical and simulated data

In order to find the best agreement between the empirical and simulated FC’s, three different metrics that capture the static as well as the dynamic organization of brain oscillations were computed across *G* (from 0 to 6 in steps of 0.1) for the mean ON, OFF and Healthy FC’s. The first one is the static *fitting* between the empirical and simulated FC matrices(Nakagawa et al. 2013) computed as Pearson correlation coefficient of the connectivity values and captures the static agreement of the underlying activity. The second metric that we used is the *Kolmogorov-Smirnov distance* (ks-d) between the empirical and simulated distribution of phase differences (see previous paragraphs) across time, which reflects dynamic instead of static properties of the network. The third and final one is the *metastability* (Deco *et al.* 2016), which reflects the overall variability of a system’s oscillations across time here derived from the standard deviation of the Kuramoto order parameter (Cabral et al. 2011):

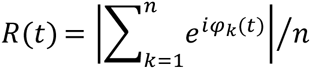

where φ_*k*_(*t*) represents the phase of all BOLD signals in a given node *k* and *n* is the number of nodes in the network. When *R* = 1 all phases are fully synchronized whereas *R* = 0 means that all phases are complete desynchronized. To do this, it is required to filter the BOLD signals with a band-pass of 0.04-0.07Hz and further compute the instantaneous phase of each narrowband signal *k* by applying a Hilbert transform in which the phase is analytically represented in:

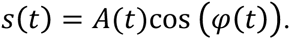

where *s(t)* is the analytic representation of a narrowband signal with an instantaneous phase *φ*(t) and amplitude *A(t).* These two elements are represented as the argument and modulus respectively in a complex signal z(t) = s(t) + *i*.H[s(t)], where *i* is the imaginary part and H[*s*(*t*)] is the Hilbert transform of *s(t).*

### Comparison of parameter distribution over conditions

To generate a distribution of bifurcation parameters for each node, we selected the optimized parameters over 20 global coupling strengths *G* in all three groups. Because we want to investigate what is the impact of DBS on global instead of local bifurcation dynamics, we explored and compared the shapes of both distributions by measuring in each condition the kurtosis *k*, which describes the shape of the distribution and the second order raw moment μ_2_, which captures data dispersion from zero. A third distribution computed from the Healthy group was also used as baseline control to see weather the overall shape is restored by the stimulation.

We used a permutation test for *k* and μ_2_ to assess recovery and a possible shift to the healthy regime. For this, we created a joint distribution from the original ON and OFF bifurcation parameter distributions. We first calculated the observed *k* and μ2 absolute difference between the ON and OFF groups which are given by:

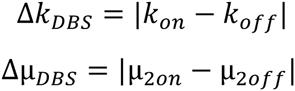

Both observed differences were compared to those of 10,000 randomized surrogate samples extracted from the joint distribution. Then, the corresponding observed differences extracted from the Healthy control and ON groups where then generated:

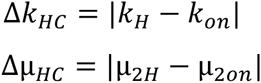

To further apply the same permutation analysis with the randomized *k* and μ_2_ versions from the joint distribution of the ON and Healthy groups.

Next, we computed the ks-d between conditions to estimate the distance between distributions. Also, for each node in the ON and OFF conditions, we computed the mean bifurcation parameter value to understand the local behaviour with respect to the global. Finally, the same semi-random walker algorithm described in previous paragraphs was applied to the modelled ON, modelled OFF and modelled Healthy networks to corroborate if diffusion of information (Rosvall and Bergstrom 2007) is also enhanced in modelled data.

### Recreating in silico local stimulation

To find which regions contribute more in shifting whole-brain dynamics to that of the healthy regime, instead of estimating all local bifurcation parameter values (see modeling paragraphs), we fixed the *a* parameter to a positive value of 0.25 (stable oscillatory regime), one node at a time in DBS OFF and repeated the modeling procedure 1000 times creating evoked bifurcation patterns from DBS OFF for each node. This method allowed, under the context of the current model, recreating stable local oscillatory conditions representative of an *in silico* stimulation. Finally, we estimated the Euclidean distance of the evoked *a* parameter vector (one value per node) from DBS OFF to that of the Healthy regime given by:

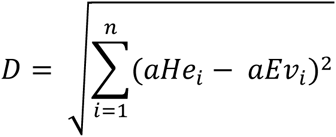

The distance *D* portrays which *in silico* stimulation site in DBS OFF brings large-scale dynamics closer to those observed in the Healthy brain. Here, *aHe* represents the Healthy bifurcation reference vector computed as the mean bifurcation parameter per node while *aEv* the evoked DBS OFF vector and *n* is the number of nodes (45 in this case). The node with a fixed parameter value was not included for all Euclidean distance estimations.

### Statistical analysis

For all analyses, a fixed p-value of 0.05 was used to determine significance. Mean metrics are always described with the mean ± standard deviation. In the case of the empirical data, the following criteria were followed. Given that DBS ON and OFF represents a paired sample, we compared the integration, phase consistency and information flow with a two-tailed paired *t*-test. To compare ON with the Healthy data set, we used a two-tailed unpaired *t*-test. Given that the distribution of modeled bifurcation parameters (Fig. 3) presented a skewed distribution not appropriate for a simple *t*-test, we employed a permutation test using 10,000 random surrogates to address statistical differences between the parameters of the distributions (see methods above). For diffusion of information, we launched the algorithm 10,000 per network to have a sufficiently large sample to compute the mean number of steps *S* and further compare these distributions with a *t*-test.

## Results

The differences in both empirical and simulated functional connectivity data for DBS OFF and DBS ON in patients and Healthy participants were measured using a number of sensitive methods. For the empirical data of each participant we measured the 1) integration, 2) mean phase consistency and 3) the standard deviation of the phase consistency (see Methods). Overall, these measurements showed significant differences between the DBS OFF, DBS ON and Healthy group, where, as expected, the highest values were found for the Healthy participants followed by lower values in patients with the DBS ON and lowest in DBS OFF (see Fig. 1). The mean phase consistency as well as the standard deviation of the phase consistency are significantly higher (p<0.0001) in the ON condition (0.267±0.027 and 0.257±0.011 respectively) compared to the OFF condition (0.191±0.041 and 0.227±0.001 respectively) suggesting that turning the stimulation ON creates more variable states, resembling that of the healthy brain (Fig. 1). Compare this to the significantly (p<0.0001) higher mean (0.402±0.053) and standard deviation (0.293±0.017) of the Phase consistency in the Healthy participants in contrast to the ON condition. This trend was also supported by a higher global integration value in the ON condition (0.621±0.089) compared to the OFF condition (0.594±0.073) (p=0.11) and even higher (0.770±0.077) and significantly different in the control Healthy group (Fig. 1).

**Fig. 1:**
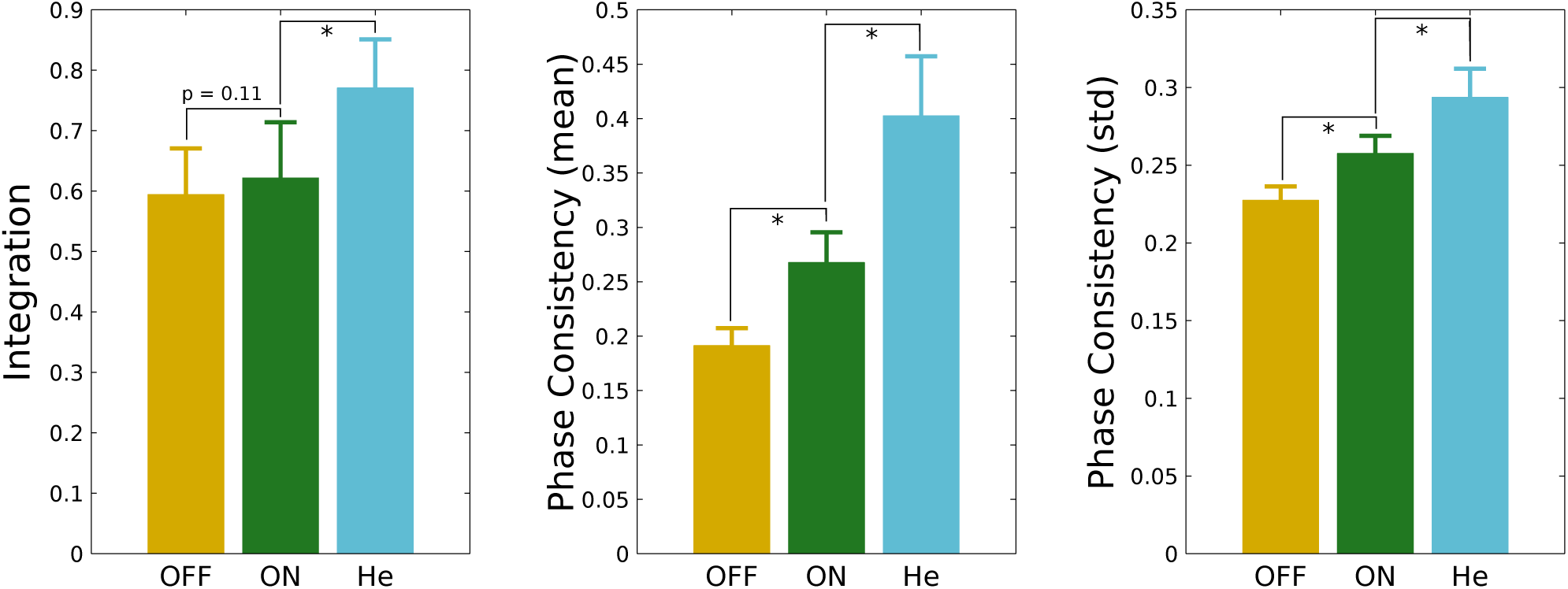
DBS induced changes in global measurements furnishing integration and metastability. These changes were seen in integration, mean phase consistency (coherence) and mean standard deviation of the phase consistency for the empirical neuroimaging datasets for the Healthy participants (He, light blue), DBS ON (ON, green) and DBS OFF (OFF, orange). As expected, the measurements are significantly highest for the Healthy group followed by the DBS ON and with lowest scores for the DBS OFF. Differences between ON and OFF correspond to a two-tailed paired i-test, while differences between ON and Healthy correspond to a two-tailed unpaired i-test. All significant differences are marked with a star and correspond to a *p* < 0.0001.

We also measured the global agreement between the empirical data and the simulated data both in a static and dynamic manner for all three groups. As shown in Fig. 2, the fitting between the simulated and empirical FC rapidly increased as a function of the coupling strength, G, and reached a plateau at around 2 for both the ON and OFF groups. Here, in accordance with the previous study by van Hartevelt and colleagues (2014) who addressed the positive shift of the global coupling required to simulate a network post-DBS in patients with Parkinson’s Disease, the maximum fit is higher in the ON (0.6) compared to the OFF (0.5) condition and even higher in the Healthy group at *G* ~ 2. As for the ks-d, it rapidly decreased in all groups reaching values of 0.1 also for *G* ~ 2 (Fig. 2), which reflects a better agreement of the dynamic properties of the network. Finally, metastability also showed a similar trend, reaching a plateau after values of *G* ~ 2 for both groups. At this coupling, the ON and Healthy groups showed a metastability value of around 0.18 while the OFF group presented a value of 0.14, suggesting that DBS enhances and restores global synchronization, even immediately after turning the stimulation on.

**Fig. 2:**
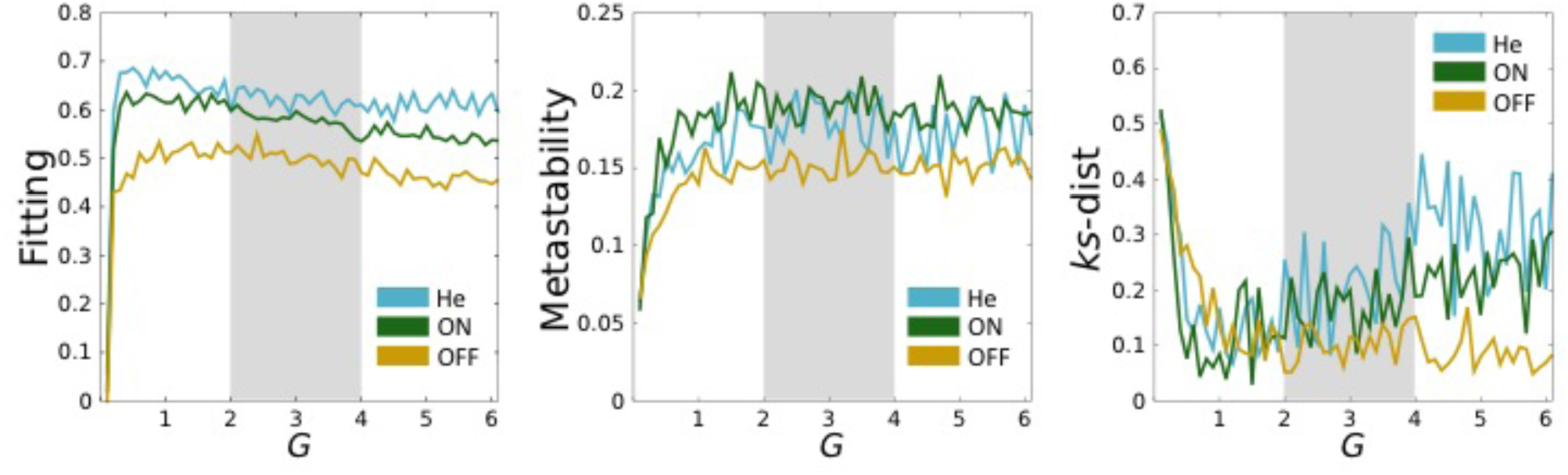
Measuring whole-brain computational modeling of DBS empirical data. The Fig. shows the agreement metrics between simulated and empirical data for all three groups of; Healthy participants (He, light blue), DBS ON (ON, green) and DBS OFF (OFF, orange). The three panels represent the measurements of fitting, metastability and Kolmogorov-Smirnov distance (see Methods) for all groups as a function of the coupling strength parameter G. The gray area represents the 20 continuous couplings from which the bifurcation parameter values where selected to construct their corresponding distributions. As expected the worst values were found for DBS OFF and significantly improve for DBS ON, almost reaching the levels of the Healthy participants.

Next, we extracted the optimized bifurcation parameters of each node from 20 coupling strength values (Fig. 2) in all groups (see Methods). Direct comparison of the distributions of bifurcation parameters between the Healthy, ON and OFF groups show that the shape of the distribution in the ON and Healthy groups is remarkably similar with a sharper peak near the bifurcation (Fig. 3). This is reflected by higher *k* and lower μ_2_ values from the ON condition compared to the OFF condition (see Fig. 3A). Investigating the Kolmogorov-Smirnov distances between parameter distributions we found that it was lowest between Healthy and DBS ON. We also thresholded the bifurcation parameter values found within the range −0.5 to 0.5 and found around 50% of the parameter values in the Healthy condition are within a threshold area of bifurcation (Fig. 3B), similar to that found for the ON condition (44%) but much lower in the OFF condition (18%). The permutation test shows that the observed differences in kurtosis (Δk_DBS_) and second order moment (Δ_μDBS_) between ON and OFF conditions are higher compared to the same differences extracted from random surrogates falling in the 99^th^ and 100^th^ percentile respectively (Fig. 4). Although a similar result can be seen for Δk_Hc_, which falls in the 99^th^ percentile compared to all surrogates, this trend is not significant for Δ_μHC_, as it is in the 61^st^ percentile suggesting a recovery towards the healthy regime.

**Fig. 3:**
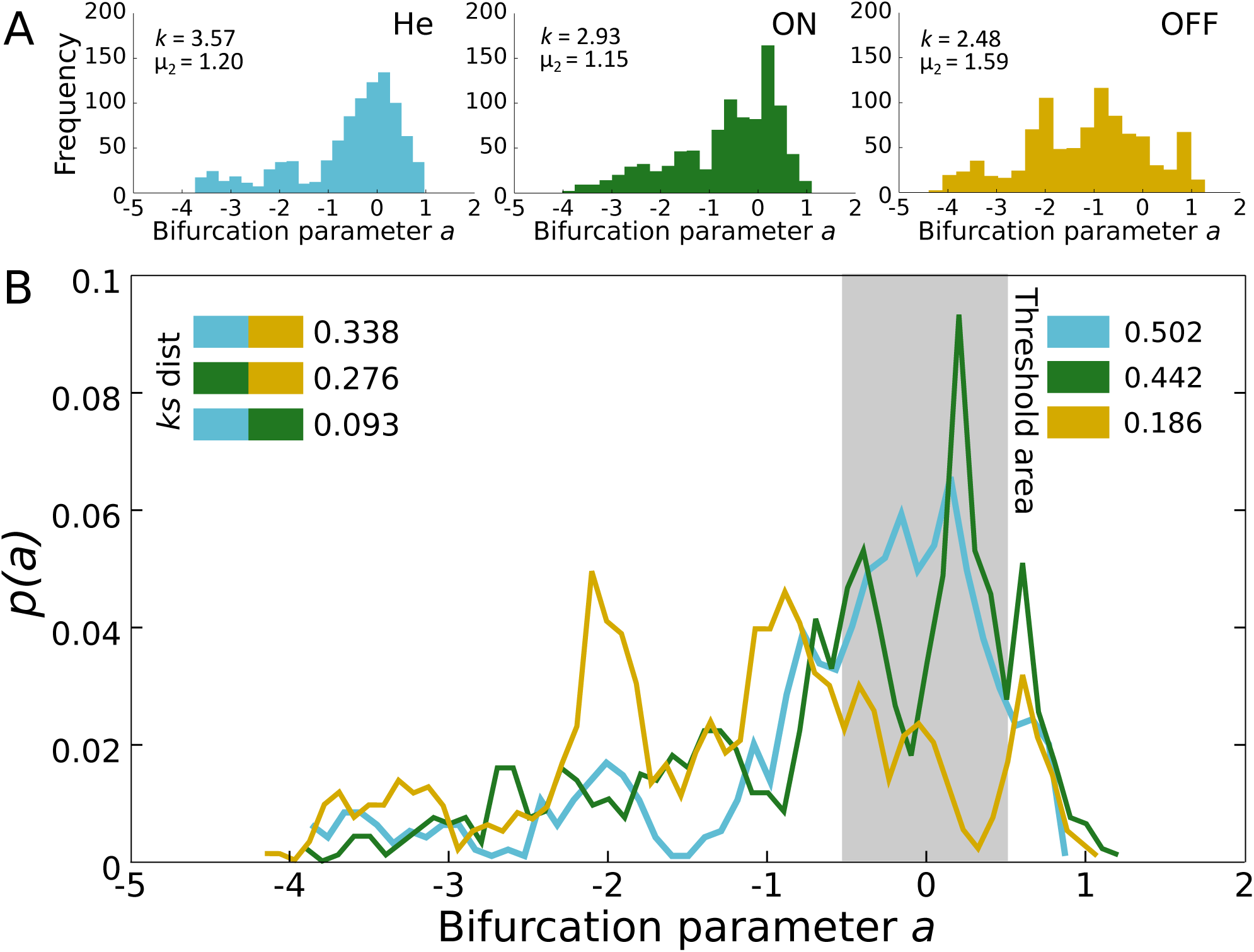
Changes in local bifurcation parameters induced by DBS. Empirical data is fitted to a Hopf model with an optimal bifurcation parameter in each local region and the resulting histograms and probability distributions are plotted for the DBS ON (ON, green), DBS OFF (OFF, orange) and Healthy participants (He, light blue) groups. As can be seen in (A) we found that the He and ON distributions were very similar while different from the OFF distributions (see the kurtosis of each distribution, k, and the second order momentum, μ_2_). Equally as shown in (B) measuring Kolmogorov-Smirnov distances we found strong similarity between the parameter distributions of He and ON (shown in the top left quadrant, with the colors representing the groups compared). The Healthy and DBS ON groups exhibited a larger probability than the DBS OFF of falling within the threshold area with the percentage of bifurcation parameter values found within the range −0.5 to 0.5 in each condition.

**Fig. 4:**
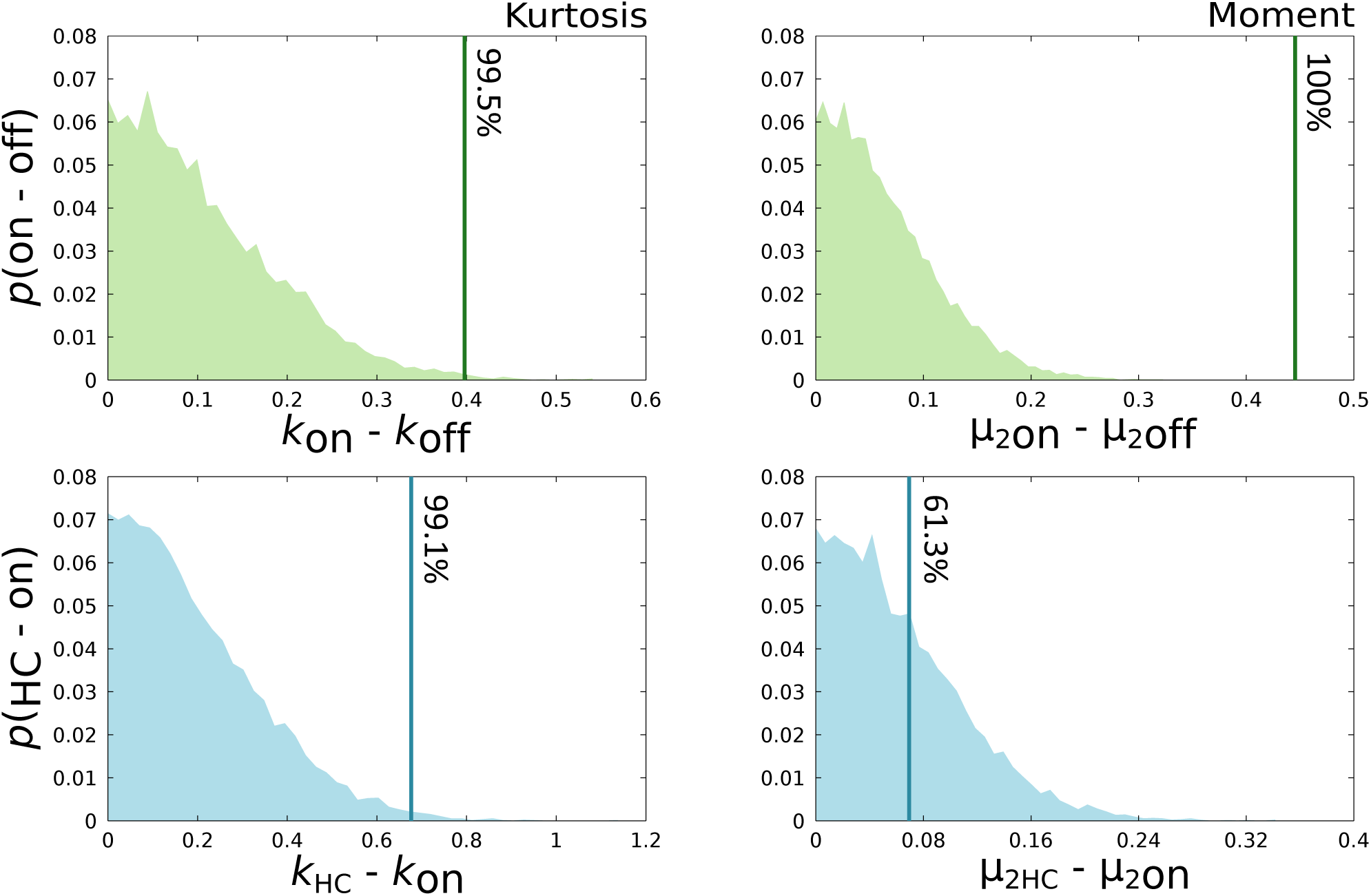
Permutation test for global bifurcation parameter distributions. The test showed significant differences between DBS ON vs DBS OFF (top row) distributions and between Healthy vs DBS ON (bottom row). The figure shows the permutation tests for kurtosis (left column) and second order moment (right column). The green histograms represent the distribution of surrogate absolute differences between DBS ON and OFF with their corresponding observed Δ*k*_DBS_ and Δμ_DBS_ differences as vertical lines and matching percentile values (see methods). The blue histograms and vertical lines represent the same analysis, but for the difference between Healthy and ON.

We also inspected the bifurcation parameter values across nodes, which allowed the identification of significant large changes between ON and OFF conditions (Fig. 5A). Regions such as the thalamus and the globus pallidus presented a big shift from highly asynchronous in the OFF condition to bifurcation values nearer criticality and ranked within the top 10 nodes with the most pronounced bifurcation parameter change (Fig. 5B & C). Interestingly, these regions are two of the main targets of DBS for Parkinson’s Disease treatment (Krause et al. 2001; Follett et al. 2010; Odekerken et al. 2013). Other regions also ranking within the top 10 nodes with the most pronounced shift are the supplementary motor area, middle cingulate gyrus as well as the insula, and the orbital part of the middle and inferior frontal gyrus switching from asynchronous in the OFF condition to values of almost 0 and even oscillatory in the ON condition (Fig. 5A & C). In contrast, only the posterior cingulate, the orbital part of the superior frontal gyrus, the triangular part of the inferior frontal gyrus and Heschl’s gyrus presented the opposite switching, from less asynchronous in the OFF to more negative in the ON condition (Fig. 5A). On the other hand, many regions such as the olfactory cortex, amygdala and hippocampus showed a higher asynchronous oscillatory behaviour in the OFF condition and slightly decreased towards a bifurcation parameter value of 0 in the ON condition suggesting that the overall distribution in the ON condition is pushing the whole brain working point closer to a bifurcation between asynchronous and stable oscillations.

**Fig. 5:**
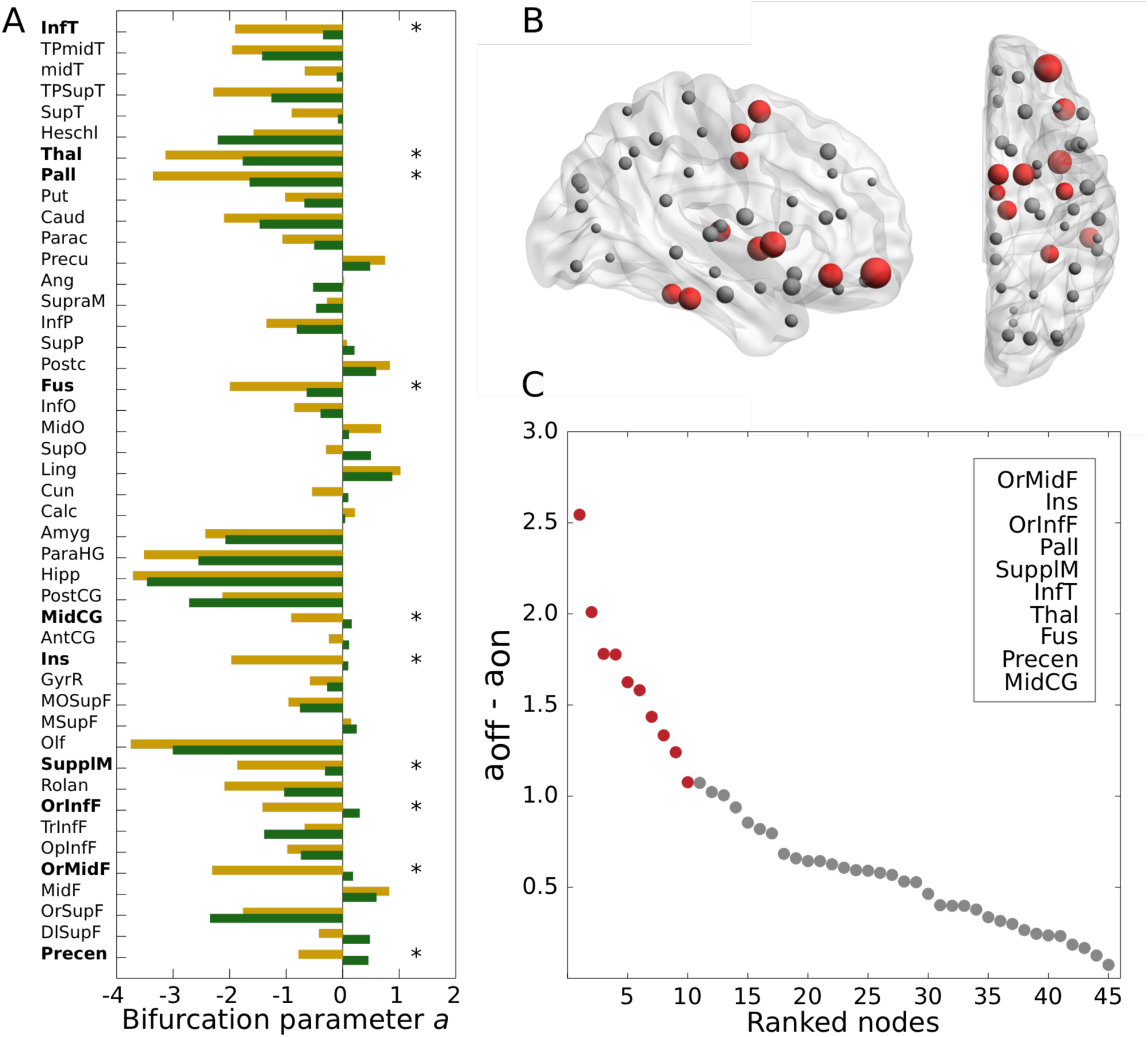
Local differences in bifurcation parameter values for Parkinson’s Disease patients. (A) Here, we show the full bar plot of the mean bifurcation parameter a in each of the 45 nodes from the right hemisphere for both the ON (green) and OFF (orange) condition. Stars and bolded regions highlight nodes with the most pronounced shift (B) Sagittal and axial view of a brain depicting the absolute bifurcation parameter shift of all nodes. Size represents shift magnitude and red nodes are those ranking in the top 10 with the largest shift. (C) Bifurcation parameter shift represented as the absolute difference between a_off_ and a_on_. Top 10 nodes are depicted in red and listed in the top-right insert. Table 1 shows the full names of abbreviated brain regions within the AAL parcellation shown. 3D brain generated with BrainNet Viewer (Xia et al. 2013).

Remarkably, our random walker algorithm showed that turning the stimulation ON creates small but highly significant improvements in diffusion of information over the network (Fig. 6), reflecting a more efficient state. In the empirical data set, the mean amount of steps *S* required for the walker to diffuse over 90% of the network is significantly (p<0.0001) lower in the DBS ON condition (117±24) compared to the DBS OFF (125±31) condition closely resembling the amount required in the Healthy group, suggesting that turning the stimulation on has a positive impact on global communicability. This effect is also present and significantly different (p<0.0001) between the modelled ON and OFF networks with mean values of 113 (±21) and 118 (±23) respectively.

**Fig. 6:**
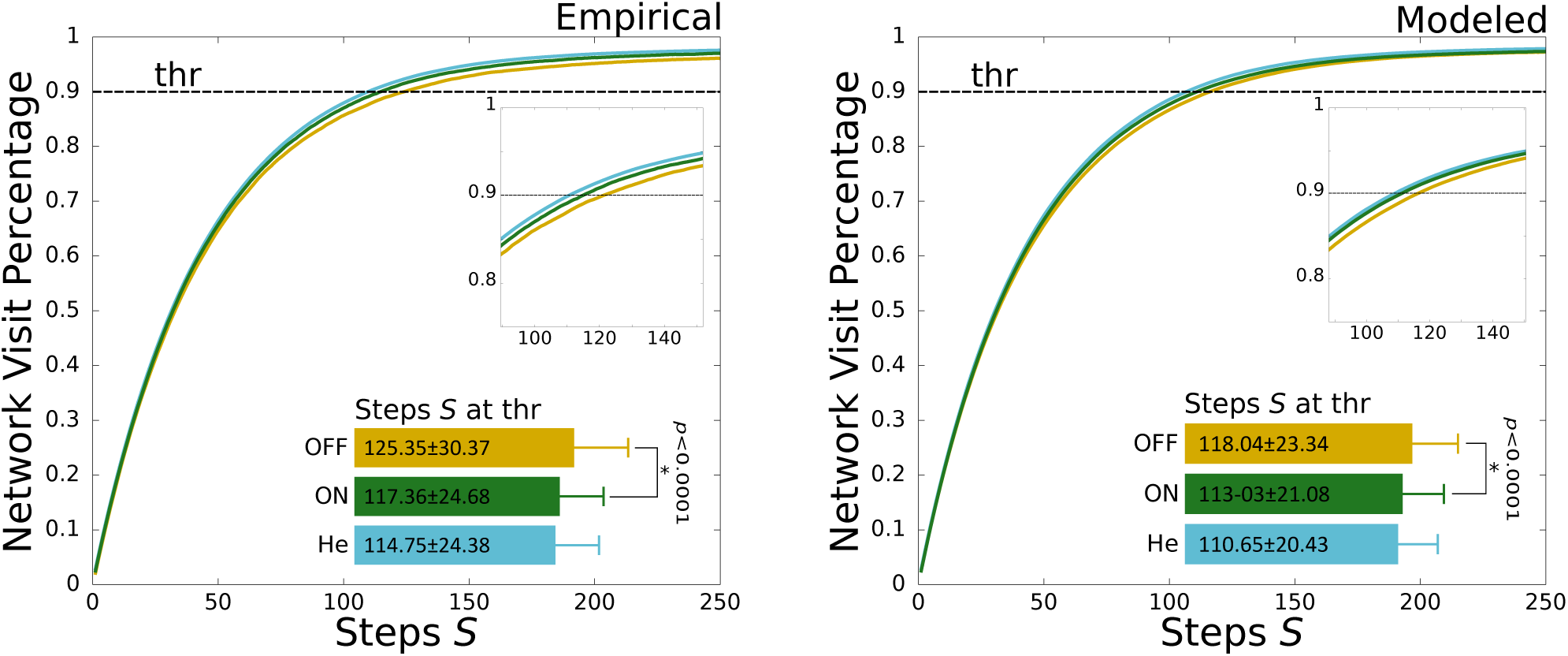
Integration capacity by using a random walker for empirical (left) and modeled (right) data. The network visit percentages of the random walker are shown for the ON (green) and, OFF (orange) conditions, as well as in the Healthy (He, light blue) group as a function of steps (S). We found that the Healthy group required the fewest steps to reach the threshold, followed by the ON group, which in turn required significantly less steps than the OFF group. Significance was assessed with a paired t-test. The horizontal dotted line represents the threshold for the number of steps *S* for which the walker has visited 90% of the network. For both data sets, bars represent the mean *S* ± standard deviation. Inserts represent zooming at threshold percentage.

Finally, our artificial *in silico* representation of local stimulation showed that some regions contribute more in shifting global dynamics of DBS OFF to that of Healthy participants as depicted by the mean Euclidean distance between the evoked (see Methods) and the Healthy reference bifurcation vector (Fig. 7). These regions are among others the putamen, caudate nucleus and the supplementary motor area (Fig. 7B), all three involved in Parkinson’s Disease therapy (Spencer et al. 1992; Montgomery et al. 2011; Niethammer et al. 2013; Shirota et al. 2013).

**Fig. 7:**
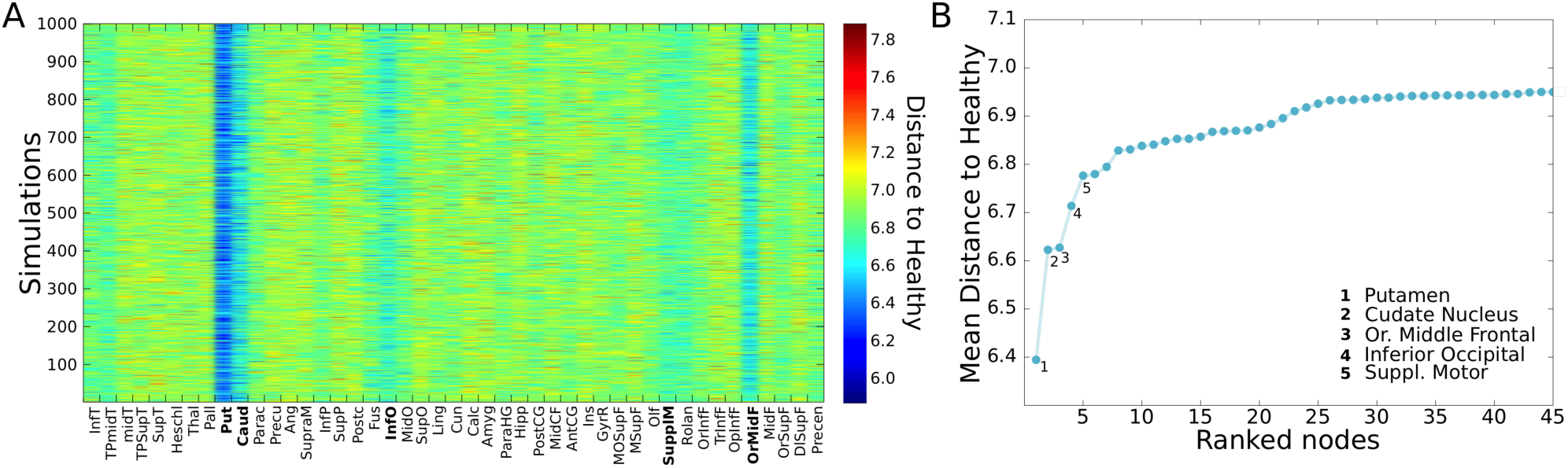
Euclidean distance to Healthy from DBS OFF after simulation of stable local oscillatory conditions. (A) Color map representing the Euclidean distance to Healthy from DBS OFF after creating stable oscillatory conditions in each of the 45 nodes. Top five regions with the lowest distance are bolded (B) Mean Euclidean distance ranked from lowest to highest.

## Discussion

The research presented here has led to novel insights into the mechanisms of DBS, using computational connectomics to model the whole-brain changes elicited by therapeutic DBS in Parkinson’s Disease. We studied the neuroimaging data from ten Parkinson’s Disease patients and found that the overall working point of the brain is shifted towards a critical, healthy state by turning DBS ON in patients with Parkinson’s Disease. More specifically, this effect is global and immediate as evidenced by better model fitting, lower Kolmogorov-Smirnov distances, higher metastability (Fig. 2) as well as higher mean phase consistency (Fig. 1) and a more efficient diffusion of the random walker in the ON condition (Fig. 5). This modeling of direct empirical observations using fMRI significantly improves on our understanding of the underlying DBS mechanisms. Among other things, this provides a better mechanistic understanding of the putative long-lasting functional impact of deep brain stimulation, previously demonstrated in a unique study of a Parkinson’s Disease patient with measurements of structural (DTI) presurgical brain changes compared to after six months of treatment (Van Hartevelt *et al.* 2014; van Hartevelt *et al.* 2015), which also showed that the working point of the brain can be partially restored by DBS.

These global changes in the working point of the brain are mirrored by specific local changes as shown by our results. We found, for example, that therapeutic DBS led to shifting of bifurcation parameter values of the thalamus and the globus pallidus from negative asynchronous to values nearer zero. It is well known that the severe degradation of the dopaminergic system causes hyperactivity in the globus pallidus, which strongly affects motor function (Dostrovsky et al. 2002). These regions are also common targets of DBS for Parkinson’s Disease and essential tremor (Odekerken *et al.* 2013) and have been shown to have significant therapeutic impact on alleviating motor symptoms (Krause *et al.* 2001; Okun et al. 2009). Further, unilateral pallidotomy studies have also shown alleviation of motor symptoms and metabolic increase measured by Position Emission Tomography in the primary motor, lateral premotor and dorsolateral prefrontal cortex (Grafton et al. 1995; Eidelberg et al. 1996). Our results suggests that one possible explanation of this important change is that Parkinson’s Disease is linked to a deleterious *global* asynchronous working point of the brain (see Fig. 3) and that therapeutic DBS can shift this state towards a more critical and efficient state with better potential for maximal exploration of the dynamic potential of the brain (Kringelbach *et al.* 2015).

Our results also show significant *local* changes in bifurcation parameters in other regions known to be affected by Parkinson’s Disease. The olfactory bulb and the temporal pole of the middle temporal gyrus were recently linked with higher structural node efficiency post-DBS (Van Hartevelt *et al.* 2014). Here, both regions presented the same trend as the thalamus and globus pallidus with less negative bifurcation values, supporting the idea of both local and global shifting from asynchronous to stable oscillations by DBS ON. Following the same tendency, the supplementary motor area changed from asynchronous to a near critical behavior, while the precentral gyrus completely switched from asynchronous to stable and both ranked within the top 10 nodes displaying the largest bifurcation parameter shift (Fig. 5C). This is potentially of interest, given that a study found that Transcranial Magnetic Stimulation (TMS) of the supplementary motor area helps alleviating motor symptoms in patients with Parkinson’s Disease (Shirota *et al.* 2013). Although a different region, in line with these findings a recent study showed that DBS successfully reduces excessive neuronal phase-locking interactions during resting state throughout the motor cortex (de Hemptinne et al. 2015). Further, the insula also presented an evident shift from asynchronous to critical behavior, which is noteworthy given its tight link with non-motor symptoms in Parkinson’s Disease (Christopher et al. 2014), as well as the previously reported bold> signal increases seen in the insula during voluntary movements under STN DBS (Kahan *et al.* 2012).

Previous studies have confirmed local synchronized oscillatory behavior in the beta frequency in the subthalamic nucleus of patients with Parkinson’s Disease that is ameliorated by therapeutic replacement of L-dopa or in the presence of therapeutic STN DBS (Eusebio et al. 2011; Litvak et al. 2011). Interestingly, our results suggest that at the local level DBS is also pushing the system towards a more asynchronous local state as seen in, for example, the posterior cingulate, Heschl’s and the orbital part of the superior frontal gyrus where the bifurcation parameters fluctuated from less to more asynchronous states. Again, this functional result fits well with structural findings, e.g. by van Hartevelt and colleagues (2014) who found that all three regions present higher nodal efficiency post-DBS. The fact that many other regions such as the precuneus, angular and middle frontal gyrus presented slightly more positive values in the OFF compared to the ON condition, but that global metrics were more positive and centered around the bifurcation (~0) suggests that whether it is changing the system from asynchronous to stable or vice-versa, DBS might enhance communicability and efficiency by pushing the system towards the bifurcation. This is supported by the random walker behavior as it diffused with greater efficiency in the ON condition (Fig. 6). The results also show that the mean and the standard deviation of the phase consistency are higher for DBS ON than DBS OFF, suggesting that activity in Parkinson’s Disease is more rigid and less variable while DBS helps creating a more flexible state (also evidenced by higher metastability), which is at the same time more similar to the healthy state. Yet, the exact mechanisms of how local bifurcation dynamics affect the global working point remain unclear and future studies should help to clarify this.

Taken together, the overall distribution shape of the local regional values of bifurcation parameters across conditions are more centered around the bifurcation in the DBS ON condition, which in turn is more similar to the Healthy regime rather than the DBS OFF (Fig. 4). Extending this finding, we also found that large-scale dynamic properties such as integration, and mean phase consistency also tend to become larger in both the empirical and simulated data sets by turning the stimulation on, suggesting that DBS has a positive effect on large-scale communicability of the network which helps rebalance the brain nearer the healthy regime (Kringelbach *et al.* 2011).

As mentioned earlier, not many studies have analyzed the impact that DBS has on the global brain activity in patients suffering from Parkinson’s Disease. Recently Kahan *et al* (2014) found that the overall effective connectivity of motor cortico-striatal and thalamo-cortical pathways is increased by DBS. Interestingly, this enhancement of connectivity strength was accompanied by reduction of clinical impairment. This fits well with the results of van Hartevelt and colleagues (2015) who found that the overall integration and segregation is improved after six months of DBS treatment in patients with Parkinson’s Disease, supporting the idea that communicability and information flow should be enhanced by the stimulation. Our random walker algorithm backs this idea, as turning the stimulation on immediately eased and restored the overall communicability of the network (measured as faster diffusion of the random walker), which was observed in both the empirical and simulated data. Owing that functional connectivity patterns are disrupted in the default mode network in patients with Parkinson’s Disease (van Eimeren *et al.* 2009; Yao et al. 2014) and that resting state effective connectivity seems to be reshaped by DBS (Kahan *et al.* 2012; Kahan *et al.* 2014), the results of the present study point towards a large-scale rebalancing as the result of specific local changes.

Recording methods that can capture activity at faster time scales such as electrocorticography (EcoG) and electroencephalography (EEG) have shown that Parkinson’s Disease is represented by hypersychronization on the beta band (8-35 Hz) in the sensorimotor network and the STN (Whitmer et al. 2012), which is interestingly restored both after DBS (Wingeier et al. 2006; Kuhn et al. 2008) and by dopaminergic therapy (Weinberger et al. 2006; Ray et al. 2008). Still, it is well recognized that DBS at the STN is able to improve motor symptoms that persists in medicated patients and that the effect is long-lasting (Kleiner-Fisman et al. 2002; Lang et al. 2003) which represents a significant improvement in quality of life compared to medicated patients (Deuschl et al. 2006). Additionally, local filed potential (LFP) recordings have shown strong synchronization in the basal ganglia in patients with Parkinson’s Disease (Brown and Williams 2005; Kuhn et al. 2006; Hammond et al. 2007). Although informative on temporal aspects, all these studies offer local information only. At a whole-brain scale, a study that used magnetoencephalography (MEG) found that network organization in Parkinson’s Disease is shifted towards a random structure representing a less efficient state (Olde Dubbelink et al. 2014). In addition, the same study found that through multiple frequency bands, node efficiency is reduced in orbitofrontal parts (Olde Dubbelink *et al.* 2014). This is interesting and in line with our findings as although locally Parkinson’s Disease seems to be represented by hypersynchrony (Brown and Williams 2005; Kuhn *et al.* 2006; Hammond *et al.* 2007), globally and on a large scale, brain dynamics in Parkinson’s Disease are less predictable and thus less efficient, also evidenced by techniques capable of recording faster oscillations compared to fMRI.

As a first step in finding novel and more efficacious targets for DBS in Parkinson’s Disease without clinical intervention and directly informed by computational models, we showed that forcing local stable oscillatory conditions in some regions pushes the system closer to the Healthy regime. Interestingly, the two regions ranking at the top are the putamen and caudate nucleus, both parts of the basal ganglia, which are clearly involved in Parkinson’s Disease (Kish et al. 2008; Montgomery *et al.* 2011; Niethammer *et al.* 2013). Even so, it is not clear if DBS in these areas would be able to alleviate symptoms. A study found that DBS in the putamen might help improve motor symptoms (Montgomery *et al.* 2011) in Parkinson’s Disease although the accuracy of the stimulation location has been questioned (Hariz 2012). Another region ranking high was the supplementary motor cortex. Notably, stimulating this region through TMS helps relieving motor symptoms in Parkinson’s Disease (Shirota *et al.* 2013) and functional connectivity between this region and the putamen is higher in patients with Parkinson’s Disease compared with controls (Yu et al. 2013), suggesting higher communicability between these two regions in Parkinson’s Disease. Other regions in the top 5 were the orbital part of the middle frontal gyrus and the inferior occipital gyrus. Interestingly, a study found that the middle frontal gyrus and the supplementary motor area are functionally implicated in visual hallucinations in patients with Parkinson’s Disease (Goetz et al. 2014) while structurally, there are differences in the middle frontal gyrus between Parkinson’s Disease with and without dementia (Goldman et al. 2014). Future analyses could study the impact of stimulating these regions in the model on restoring balance in the brain.

Given that improvements in information diffusion and phase consistency were immediate, it would be interesting to see if these effects are long-lasting and if the bifurcation shifts towards the healthy regime also relates with better behavioral scores. Deep brain stimulation has proven to have positive results after months or even years of treatment (Rodriguez-Oroz et al. 2005; Ostergaard and Aa Sunde 2006) reshaping the structural network architecture post-stimulation (Van Hartevelt *et al.* as well as effective connectivity patterns (Kahan *et al.* 2014). It would be important then to understand if the boost in communicability is also maintained throughout the treatment both at a group level and participant level. We have shown here that DBS pushes the overall dynamics of Parkinson’s Disease patients towards the healthy regime. This is consistent with a recent study showing default mode network functional connectivity differences in Parkinson’s Disease patients compared to controls (Yao *et al.* 2014). It should be noted that future work should try to control the confound potentially introduced by the resting hand tremor in patients when DBS is turned off. Furthermore, it will be important to replicate the findings using control participants scanned on the same scanner. Equally, future studies could potentially clarify if global changes caused by DBS result in more similar communicability patterns compared to healthy controls which are not only immediate but long lasting and are accompanied with cognitive and motor improvements in patients without irreversible neurodegeneration such as the dopamine depletion in Parkinson’s Disease but e.g. patients with chronic pain or other autonomic effects (Hyam et al. 2012). As such computational models might be able to inform the development of novel, more efficacious DBS targets, although it is important to carefully consider the ethical challenges of DBS (Kringelbach and Aziz 2009, 2011).

In this study, we have explored the impact that therapeutic deep brain stimulation has on global human whole-brain dynamics. More specifically, by using a neuro-dynamical model we were able to identify both local and large-scale changes necessary for the optimal fitting of the model to DBS OFF, DBS ON and Healthy participants. Remarkably, we were able to show that DBS shifts the overall brain dynamics towards the bifurcation (rather than towards noisy or asynchronous oscillatory states) and thus closer to the dynamical regime found in the healthy brain. This finding is further supported by our findings of an enhancement in global synchrony and communicability, as reflected by a higher mean phase coherence and more efficient diffusion of information in the DBS ON condition, again pushing the system towards the healthy regime in both empirical and simulated data. We also showed that forcing local stable oscillatory conditions in some regions as a proxy for stimulation pushes the system closer to the Healthy regime, especially at the putamen and globus pallidus. Finally, we showed that global changes in extrinsic connectivity were associated with local changes in intrinsic connectivity (bifurcation parameters) in very specific brain regions. Future studies are required to further clarify the mechanisms underlying these local leading to global enhancement and whether there may in fact be other DBS targets that can better rebalance the brain dynamics back to a healthy state.

## Funding

In this work, Gustavo Deco is supported by the European Research Council (ERC) Advanced Grant DYSTRUCTURE (n. 295129). Morten Kringelbach is supported by the ERC Consolidator Grant CAREGIVING (n.615539) and the Center for Music in the Brain, funded by the Danish National Research Foundation (DNRF117). Victor M Saenger is supported by the Research Personnel Training program PSI2013-42091-P funded by the Spanish Ministry of Economy and Competitiveness.

